# Inferring the T-cells repertoire dynamics of healthy individuals

**DOI:** 10.1101/2022.05.01.490247

**Authors:** Meriem Bensouda Koraichi, Silvia Ferri, Aleksandra M Walczak, Thierry Mora

## Abstract

The adaptive immune system is a diverse ecosystem that responds to pathogens by selecting cells with specific receptors. While clonal expansion in response to particular immune challenges has been extensively studied, we do not know the neutral dynamics that drive the immune system in absence of strong stimuli. Here we learn the parameters that underlie the clonal dynamics of the T-cell repertoire in healthy individuals of different ages, by applying Bayesian inference to longitudinal immune repertoire sequencing (RepSeq) data. Quantifying the experimental noise accurately for a given RepSeq technique allows us to disentangle real changes in clonal frequencies from noise. We find that the data are consistent with clone sizes following a geometric Brownian motion, and show that its predicted steady state is in quantitative agreement with the observed power-law behaviour of the clone-size distribution. The inferred turnover time scale of the repertoire increases substantially with patient age, and depends on the clone size in some individuals.

## I. INTRODUCTION

The adaptive immune system protects us from many infections including those caused by pathogenic challenges that did not exist when we were born. This amazing plasticity is encoded, in part, in a diverse repertoire of T cells carrying surface receptors capable of recognizing different antigens, which trigger an immune response. About 10^8^ new T cells are estimated to be generated and enter the periphery in human adults every day [1, 2], where they undergo specific proliferation due to antigen stimulation but also non-specific divisions [3, 4] and death. These processes together result in clone sizes of different T cells that differ over a few orders of magnitude, forming long tailed distributions [5, 6]. The total number of different T cell clones is estimated between 10^8^ and 10^10^ [7–9]. Qualitatively describing the T cell clonal dynamics in the periphery is important for predicting long-as well as short-term immune response and to understand the maintenance of immune memory.

A lot of effort has been put into describing antigen specific response and memory formation [10–12]. At any given timepoint the majority of the T cell repertoire is not always directly involved in fighting the current antigenic challenge. Yet, processes such as homeostasis [3] and unspecific signals in both naive and memory subrepertoires result in frequency changes of background clones. Many first-principles models of naive T cells dynamics have been proposed to study the balance between thymic output and peripheral proliferation and death [2, 4, 13– 15]. The role of competition for antigens between T cells has been pointed out [16], as well the effect of cross-reactivity [17] (the ability for one T cell to recognize different antigens), the relative size of a primary versus secondary response to similar antigens [18], or the effect of beneficial mutations [19]. These studies highlight the importance for the naive repertoire of clonal expansions that are not necessarily linked to specific challenges. While these models were instrumental in advancing our understanding of bulk repertoire dynamics, and allowed for the interpretation of deutered water and bromium staining experiments that describe cellular lifetimes [20], the class of models that are consistent with the data is still large and unexplored.

Thanks to advances in immune repertoire sequencing (RepSeq) [21–23], dynamical models can now be assessed directly against repertoire data at the clonal level. RepSeq experiments isolate and sequence the T cell receptors (TCR) in a blood sample of individuals. By counting reads with the same TCR sequence, one can estimate the frequency of the corresponding clone (defined as the set of cells with the same receptor) in blood. Even single repertoire snapshots can be informative about the dynamics: the distribution of clone sizes follows a power law [6, 24–27], in accord with proposed models of stochastic growth and death [5]. Taking samples from the same individual at different timepoints allows for tracking the evolution of TCR clone sizes in time. The longitudinal experiments that have been performed in healthy donors [28, 29] suggest that the repertoire is relatively stable over years.

Our main goal in this article is to characterize the dynamics of the unstimulated background repertoire. We use an inverse approach to learn models of stochastic TCR clonal dynamics directly from data, collecting human TCR RepSeq datasets where we could identify at least two time points between which there was no reported specific acute antigenic stimulation [28–32]. A key aspect of our method is the treatment of experimental noise, which confounds naive analyses of stochastic time traces. The method first quantifies both the sampling and natural biological noise thanks to replicate RepSeq experiments [33, 34], and then infers the parameters of a stochastic dynamical model to describe the trajectories of each TCR clone population in a healthy individual. We explicitly show how correcting for noise allows us to robustly learn the underlying dynamics.

A recent study [35] has investigated the formation of the T cell receptor during development and its maintenance into adulthood. Here we focus on healthy adult repertoires that are already shaped during the first years of an individual’s life, and ask how they evolve and get renewed. We extract clone turnover time scales, and describe how these time scales depend on the person’s age. Characterizing this baseline dynamics is an important step towards interpreting TCR dynamics in the presence of antigenic stimuli.

## II. RESULTS

### A. Longitudinal sampling of TCR repertoires of healthy individuals

T-cell repertoires are large ecosystems in which each species is a clone of T cells carrying the same TCR *i* formed by a unique pair of *α* and *β* chains. The dynamics of this system is characterized by the time course of the number of cells carrying each receptor, *n*_*i*_(*t*). This number can be accessed indirectly through TCR repertoire sequencing (RepSeq), obtained by sequencing the TCR of small samples of peripheral blood mononuclear cells (PBMC), giving us a read count 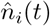 for a given chain at different timepoints (Fig. 1A). Because the two chains are not paired in the data, from here on we define clones as collections of cells having the same *α* or *β* chain, which we will refer to as clonotypes. This approximation is justified by the low occurence of TCRs that share one chain but not the other [32].

**FIG. 1:**
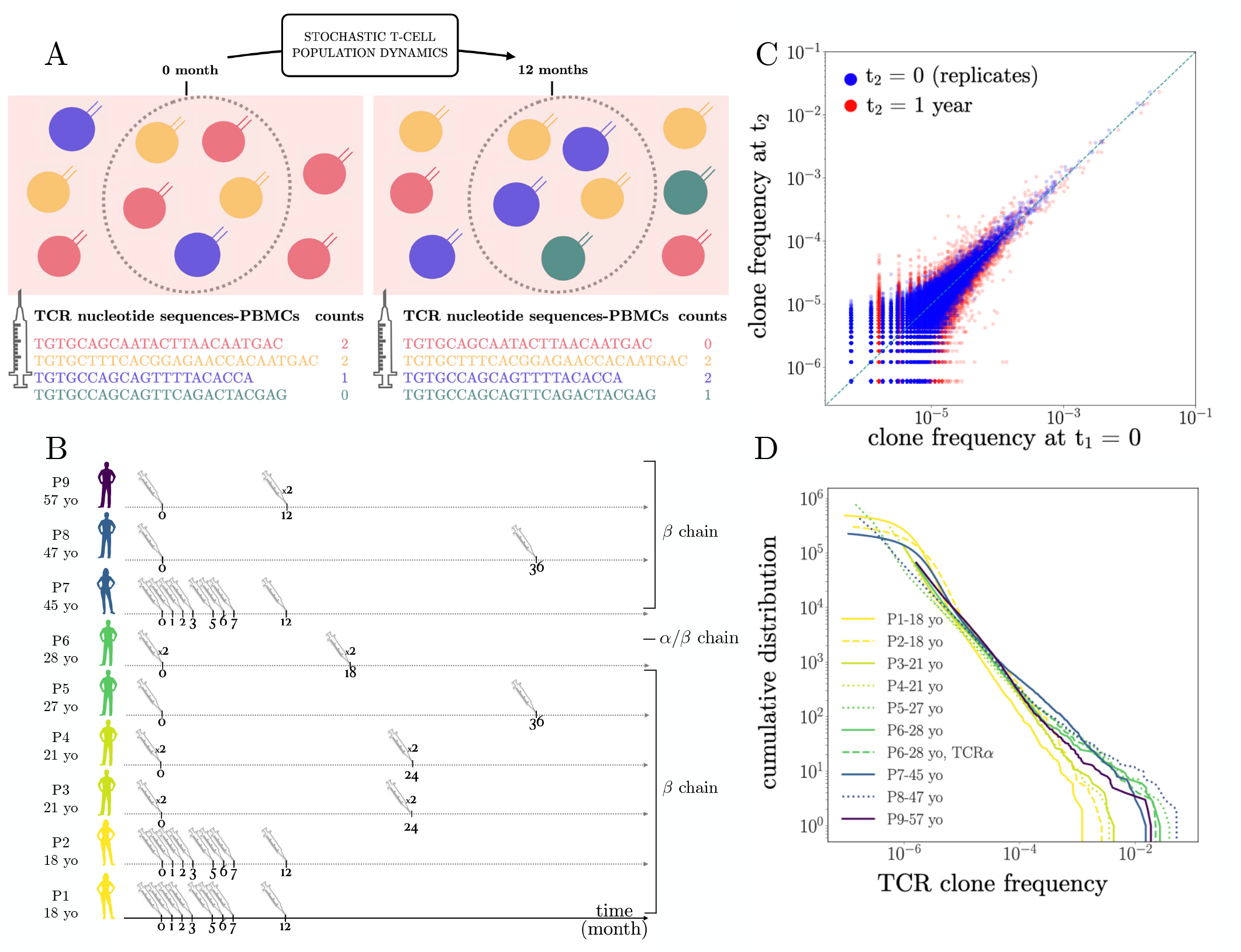
Longituinal tracking of T-cell repertoires. **A**. Experimental workflow. PBMC from a healthy individual are extracted at two timepoints, and their TCR repertoire sequenced, yielding lists of clonotypes with count numbers corresponding to the number of individual measurements (or reads). The way in which the two sampled repertoires has changed between the two timepoints is predicted by a stochastic model of the dynamics of T cell clones. **B**. Summary of the TCR *α* and *β* repertoire data used in this study. 9 individuals from 5 studies, aged 18-57, male and female, were included. When available, replicate experiments are annotated with *×*2. Datasets were produced using two different sequencing technologies based on cDNA and gDNA. **C**. Typical scatter plot of frequencies of TCR clones in two samples from the same individual P9. Blue: two biological replicates obtained on the same day show the effect of experimental nosie. Red: 2 samples taken 1 year apart show a larger spread, resulting from a combination of real changes and noise. The goal of the analysis is to disentangle real changes from the noise. **D**. Cumulative distributions of TCR frequencies, which follow a universal power law in all samples and donors, with exponent ≈ 1.

We collected repertoire data from 9 individuals P1-P9, aged 18-57, sampled at various time points from one month up to 3 years apart, with and without biological replicates. *β* chain repertoires were sequenced for all samples, and *α* chains only for individual P6. The properties of the datasets, including their number of clones *N*_*c*_, total read counts 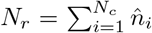, age, library preparation (from genomic DNA or from mRNA), and chain, are summarized in Fig. 1B and in Table S1. Because P3, P4, P6, and P9 were included in a vaccination study, they had received a shot of the YFV 17D yellow fever vaccine (P3, P4, P6) or of the influenza vaccine (P9) 45 days prior to the first time point, after the decay of their T-cell response, so we assume that the dynamics of vaccine-specific T does not affect much our analysis of the global repertoire.

A major challenge when analyzing RepSeq data is that the measured abundances 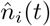 only provide a noisy reflection of the true ones *n*_*i*_(*t*). Observed differences between datasets thus result from a combination of the repertoire dynamics and biological and experimental noise. The magnitude of that noise can be assessed by comparing the normalized clonotype frequencies 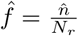 between two biological replicates obtained at the same time point in the same individual (Fig. 1C, blue dots). By contrast, comparing those frequencies between two time-points separated by one year (Fig. 1C, red dots) show a larger dispersion, and a slight overall decrease of clonotype frequencies. Our goal is to measure this difference quantitatively.

Another difficulty arises from the observation that clonotype frequencies are highly heterogeneous, with their distribution following a power law 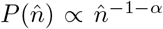spanning no less than 4 orders of magnitude, with an exponent *α* ≈1 which is largely invariant across individuals and timepoints (Fig. 1D), as previously reported [5, 6, 35]. This implies that most clonotypes have very low abundance and are thus particularly subject to sampling and experimental noise.

### B. Mathematical model of stochastic clonal dynamics

The dynamics of T cell clones is driven by the proliferation and death of cells belonging to them. In addition, new clones with their distinct TCR are continually produced and released by the thymus, although the rate of thymic exports decays rapidly with age [1]. Cell division, death, and introduction of new clones constitute the basis of our model (Fig. 2A). Cell division may be caused by antigen stimulation (both self and foreign) or by cytokine and growth factors, and cells die by lack of stimulation or by apoptotic signals. Even in the absence of strong and chronic antigenic stimuli, T cell clonotype abundances display stochastic trajectories due to either weak stimulation, repertoire homeostasis and demographic fluctuations. In addition, individuals may get mild infections over the course of months and years. Since these events are numerous and unknown, we model them by an effectively random net growth rate (divisions minus deaths). It can be shown [5, 36] that on time scales much longer than the typical resolution time of infections, each clonotype size *n*_*i*_(*t*) may then be modeled by a geometric Brownian motion (GBM). Its evolution is governed by an effective mean net growth, to which random fluctuations are added to account for bursts of proliferation and decay:

**FIG. 2:**
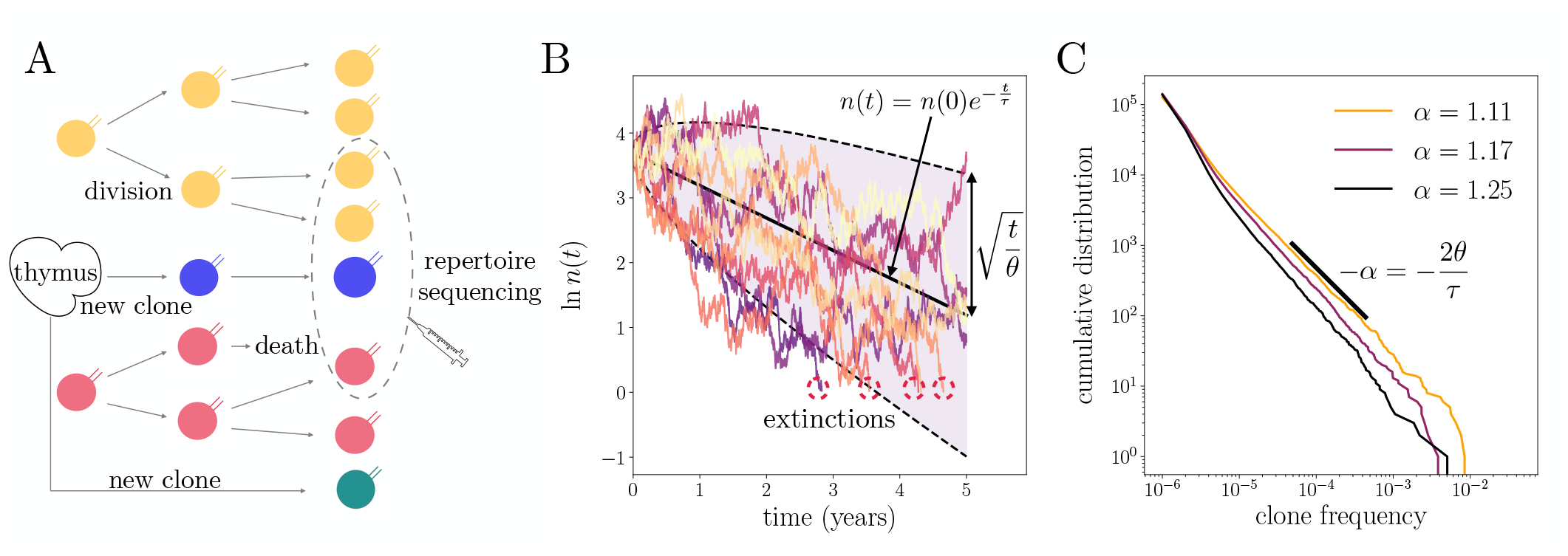
Stochastic model of repertoire dynamics. **A**. T cells are introduced in the peripheral immune system by thymic export, providing a source of new TCR clones. T cells belonging to a specific TCR clone (labeled by their color) divide and die depending on their interactions with the antigenic environment, increasing or reducing the abundance of its TCR in the repertoire. This process is modeled by a geometric Brownian motion **B**. Example traces of TCR abundances simulated from the model Eq. 1 with *n*(0) = 40, with *τ* = 2 years and *θ* = 1.11 year. Clones that reach abundance *<* 1 go extinct (red circles). The typical trend is for clones to decay exponentially with time scale *τ* (black solid line). Stochastic events of growth and decay account for a broad variability of individual traces, whose magnitude grows as 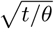 with time (shaded area) in logarithmic scale. **C**. Cumulative frequencies distributions of synthetic TCR clone abundances. The model predicts a power law of exponent *α* = 2*θ/τ*. Different values of *τ* and *θ* were used to lead to different values of the exponent *α*. Parameters: *τ* = 2 years, *N*_cell_ = 10^10^, *n*_0_ = 40.

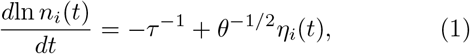

where *η*_*i*_(*t*) is a clonotype-specific white noise of zero mean and unit amplitude. Note that the mean growth rate of clones, −*τ* ^−1^, is typically negative. On average, each clone should decay to make room for new thymic exports, because of homeostatic pressures that control the total number of cells. In this interpretation, *τ* is the typical decay and turnover time of each clone, which would evolve with time as *n*_*i*_(*t*) = *n*_*i*_(0)*e*^−*t/τ*^ in the absence of fluctuations. But recall that this is just an average— many clones do not decay, but instead undergo episodes of large growth and decay, as illustrated by simulations of (1) in Fig. 2B. The typical amplitude of these fluctuations grows with time as 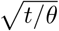 (dashed lines). Thus, *θ* may be interpreted as the typical time it takes for a clone to rise or decay above or below the typical behaviour by one log-unit.

In addition to being biological motivated, the proposed dynamics have the desirable property that, in the presence of a constant rate of thymic exports, the distribution of clone sizes is predicted to evolve in time towards a perfect power law, *P* (*n*) ∝*n*^−1 −*α*^, with exponent *α* = 2*θ/τ* given by twice the ratio of the two time scales of the model [5]. This is illustrated in Fig. 2C on simulated repertoires at steady state, and agrees well with the empirical distributions of Fig. 1D.

Our goal is to capture the parameters of these dynamics that is informative about the repertoire turnover timescales, while constraining the experimentally observed clone size frequency distribution. Our approach assumes that on the timescales of the analysis we do not observe signals of strong and specific antigenic stimulation. It also ignores potential dependences on the size of the clone, which could be mediated by phenotypic differences between clones. This last assumption will be revisited later.

### C. Model inference

We estimate the parameters (*θ, τ*) of the dynamics in Eq. (1) from the observed clonotype abundance trajectories using a Bayesian approach for the posterior distribution of parameters given the data:

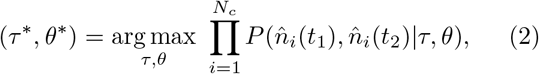

where *t*_1_ and *t*_2_ are the times of the two samples.

We use two methods to learn the model parameters: *naive inference* and *full inference*. The naive inference assumes the empirical abundances faithfully represents the real clonal abundances *n*_*i*_ through a simple proportionality rule, 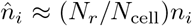, where *N*_cell_ = ∑_*i*_ *n*_*i*_ is the total number of T cells in the body. In practice, we work with clonotype frequencies 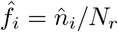, and *f*_*i*_ = *n*_*i*_/∑*N*_cell_, so that this assumption becomes 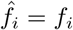. Further assuming that the total number of cells *N*_cell_ is approximately constant in time at steady state, 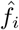 is then governed by the same equation (1) as *n*_*i*_. We take advantage of the closed solution available for the propagator associated to the GBM, (4), to maximize the log-likelihood (see Methods). This maximization is equivalent to plotting a histogram of the change in log-frequencies between the two timepoints, and simply read off *τ* ^−1^ and *θ*^−1^ as the negative mean and the variance of the distribution divided by *t* = *t*_2_ − *t*_1_ (Fig. 3A), consistent with their biological interpretation.

**FIG. 3:**
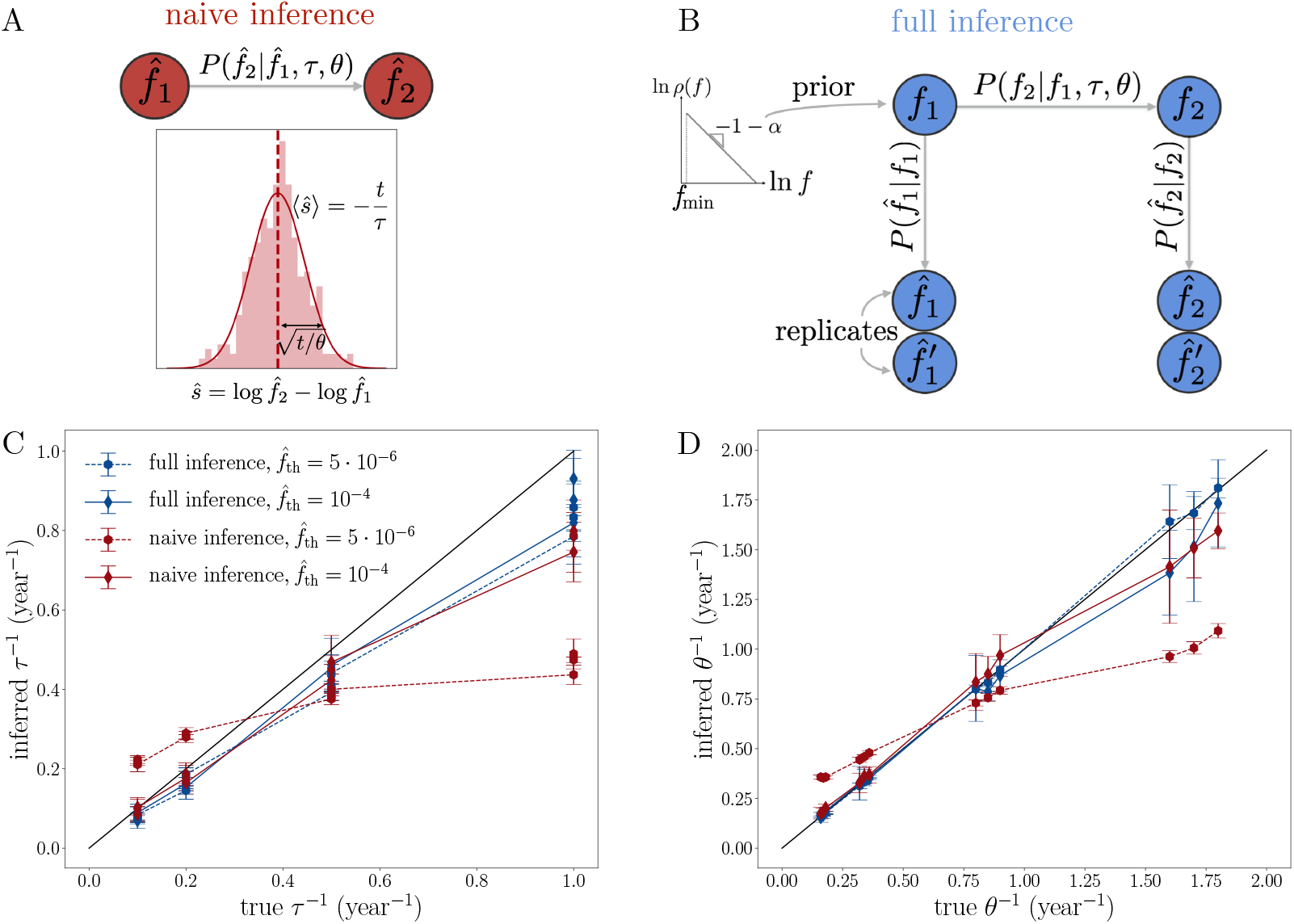
Inferring dynamical parameters from data. **A**. Naive inference. The empirical clonotype frequencies in the RepSeq samples at the two time points, 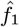 and 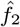, are treated as the true ones, *f*_1_ and *f*_2_. We estimate the two parameters *τ* and *θ* from the average and standard deviation over all observed TCR clones of the log-fold change in frequency between two-time points, 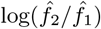, which the model predicts is distributed normally. The distribution of *Ŝ*is shown in light red, and the Gaussian fit in solid red. **B**. Full inference. The empirical frequencies 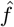 are modeled as noisy read-outs of the true ones *f*, through a probabilistic noise model. First, the noise model 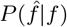 is inferred from replicate experiments such as shown in Fig. 1C. The inference procedure also learns the distribution of frequencies *ρ*(*f*), assumed to follow a power law with adjustable exponent *α* and minimal frequency *f*_min_. Second, using the noise model, the parameters of the dynamical propagator *P* (*f*_2_ |*f*_1_, *τ, θ*) are inferred from two timepoints, where *f*_1_ and *f*_2_ are treated as latent variables and 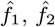 as observables, using a Maximum likelihood estimator. **C-D**. Validation of naive and full inference models on synthetic data. Model parameters: *t*_2_ − *t*_1_ = 2 years, *τ* = 1, 2, 5, 10 years, *α* = 1.11, 1.18, 1.25, with all 12 combinations tested; number of cells *N*_*c*_ = 10^10^; initial clone size *n*_0_ = 40; the other parameters (number of clones, thymic output) are deduced assuming steady state (see Methods). Sampling model: number of sampled reads *N*_*r*_ = 10^6^; noise model parameters *a* = 0.7 and *b* = 1.1. Error bars are standard deviations over 10 simulations.

The *full inference* incorporates the fact that the observed clonotype abundances are contaminated by biological (mRNA expression) and experimental noise sources (sequencing errors, stochastic PCR amplification, and sampling), which means they do not correspond exactly to the clonotype abundances. To give a sense of just the sampling issue, a PBMC sample of ∼1 mL contains about 1 million cells, yielding about 1 million reads. By comparison, the organism contains of the order of 10^11^ T cells. TCR clonotype frequencies are thus extrapolated from observing a fraction 10^6^*/*10^11^ ≈10^−5^, or 0.001%, of the whole repertoire [9]. In addition, not all cells are captured, and each cell may be represented by multiple reads, either through sequencing of multiple mRNA from the same cell, or from PCR amplication, depending on the context. To address these sources of uncertainty, in the full inference approach we introduce an error model [33] relating observed frequencies 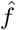 to their true value *f* probabilistically through the transfer function 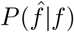 (Fig. 3B). We use the previously introduced a software tool, NoisET [34], which learns such a noise model from replicate RepSeq experiments (see Methods).

We applied NoisET to individuals P3, P4, P6 and P9 for whom replicates were available. The noise model assumes that the read count 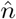 of each clone is drawn from a negative binomial distribution, whose variance grows with the frequency as 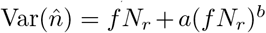, with two learnable parameters *a, b*. In addition, since true frequencies are unknown, we assume as a prior that frequencies are distributed according to a power law *ρ*(*f*) ∝*f* ^−1 −*α*^ with a cut-off *f > f*_min_, with *α* and *f*_min_ another two parameters. These parameters are reported for all individuals and time points in Fig. S1.

Once the noise model has been learned using NoisET, the likelihood of the data is computed by summing over the latent variables *f*_1_ = *f*_*i*_(*t*_1_) and *f*_2_ = *f*_*i*_(*t*_2_):

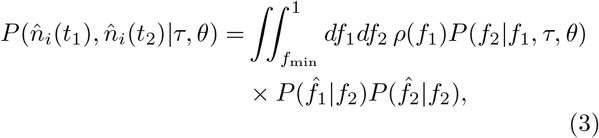

where *P* (*f*_2_|*f*_1_; *τ, θ*) is the propagator of the geometric Brownian motion Eq. 1, and 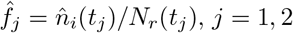.

To explore the dependence of the *τ* and *θ* parameters on the frequency of clonotypes, and to eliminate clones that are not seen at both timepoints, we can generalize the formulas above to include only clonotypes with frequencies larger than a specific threshold 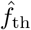, which modifies the normalization of the maximum likelihood estimator (see Methods).

### D. Validation of the inference methods on synthetic data

We first test the naive and full model inference on simulated RepSeq samples. We simulate 10^10^ cells corresponding to ∼10^8^ synthetic longitudinal trajectories designed to mimic as closely as possible the features of the real repertoire data at time points two years apart. The initial size *n*_*i*_(*t*_1_) of each clone is drawn from the steady state distribution of the GBM with a constant source (see Methods). Then Eq. 1 is simulated between times *t*_1_ and *t*_2_ with an extinction condition when *n*_*i*_ *<* 1, and with a source of new clones whose rate of introcution is matched to the mean extinction rate (see Methods). We varied the two timescales of the model, *τ* and *θ*, from months to years, while keeping *α* = 2*θ/τ* within the observed experimental range 1.1 – 1.25 [35].

We model experimental sampling using a negative binomial distribution with variance parameters *a* = 0.7 and *b* = 1.1. Sequencing depth was set to *N*_*r*_ = 10^6^ reads at both time points (we checked that asymmetric numbers of reads at each timepoint did not affect the results, see Fig. S2), resulting in ∼10^5^ sampled distinct clonotypes. For each set of parameters we generated 10 longitudinal datasets to assess errors. We then performed the naive and full inference methods on these datasets, restricted to clones with 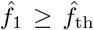 and 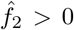, and compared the inferred values of *τ* and *θ* to the true ones (Fig. 3C-D).

While the full inference (blue points) works for all values of the parameters and frequency threshold, the naive inference (red points) performs poorly for large values of the parameters. Increasing the cutoff frequency to *f*_th_ = 10^−4^ improves the naive inference by limiting the effect of the sampling noise, which is relatively smaller in large clones. For lower values of the threshold, the more numerous small clones dominate the inference, yielding an erroneous estimate. However, since the naive inference does not require replicates or a noise model, and is faster to implement, it provides a practical solution for learning *τ* and *θ* for large clones.

### E. Analysis of repertoires

We applied the full inference to longitudinal data sets of healthy individual TCR repertoires presented in Fig. 1 for which replicates were available, focusing on large enough clones 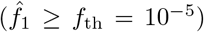. With this cutoff we limit experimental noise and focus mainly on memory clones, since large clones are more likely to be have arisen from expansion and belong to the memory pool [14]. For all individuals, the inferred values of *τ* and *θ* fall close to a line defined by *τ* ≈ 2*θ* (Fig. 4A) corresponding to a predicted exponent of *α* ≈2*θ/τ* = 1 in the power law of the clone size distribution. This result is in agreement with empirical observations of Fig. 1D. A more refined comparison of the predicted exponent, 2*θ/τ*, with the one directly inferred from the distribution of clone sizes, *α*, gives consistent but noisy results (Fig. 4B), primarily because of the narrow range of values of *α* (0.9 – 1.2) and the small number of individuals. We note that the two inferred values of *α* use completely independent pieces of information, namely the clone size distribution in one case, and the dynamics of clone sizes in the other.

**FIG. 4:**
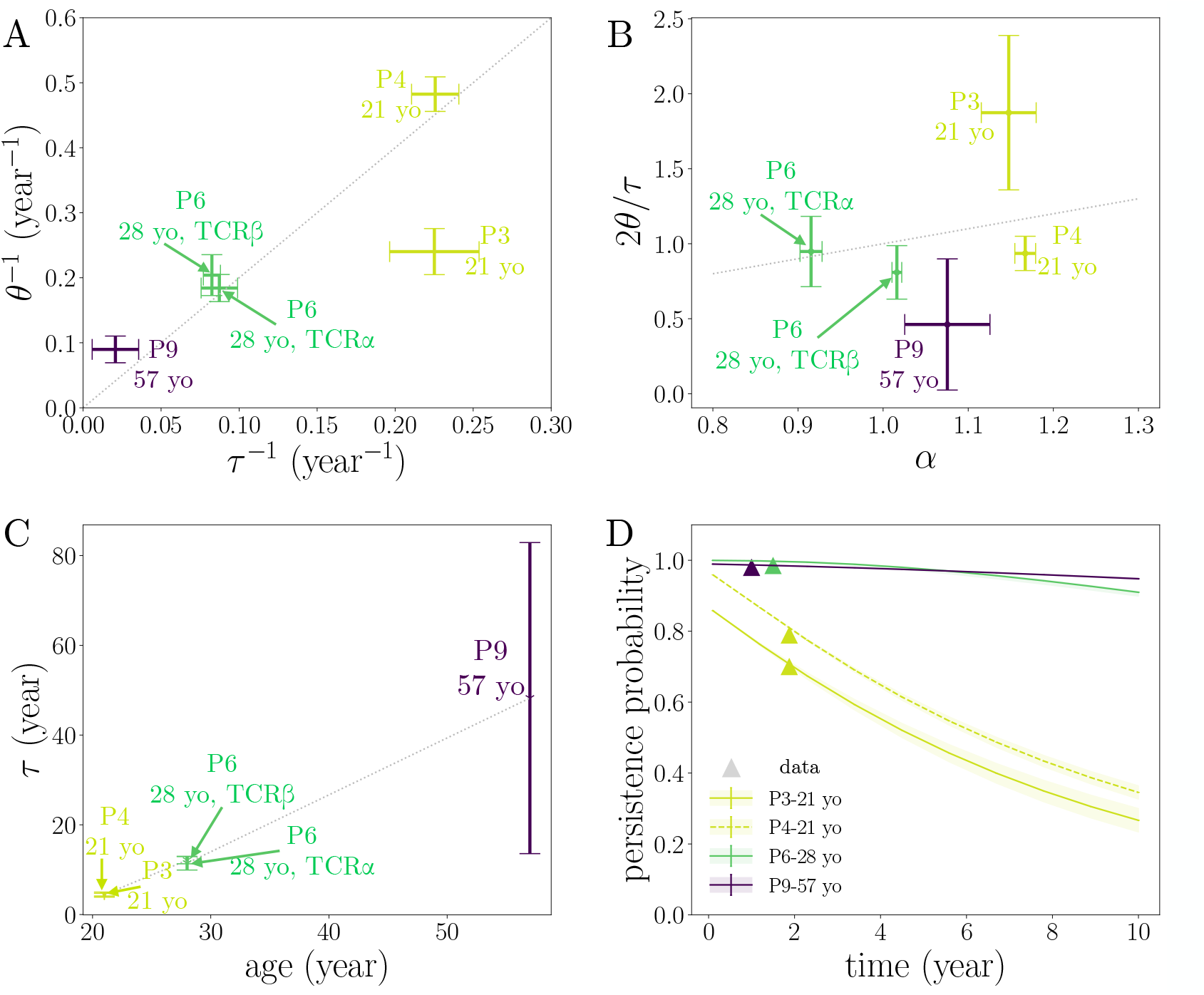
Dynamical parameters of healthy TCR repertoires. **A**. Typical decay rate *τ* ^−1^, and inverse fluctuation amplitude *θ*^−1^ for the 5 donors for whom replicates were available, as obtained using the full inference procedure with *f*_th_ = 10^−5^. All donors but one are consistent with the relation 2*θ* = *τ*, corresponding to *α* = 1. Error bars are standard deviations over all combinations of the replicates at each time point. **B**. Direct test of the prediction *α* = 2*θ/τ*. Most values of *α* fall close to one, allowing for only a narrow range of tested values. Error bars on *α* show standard deviations across time points. **C**. Turnover parameter *τ* as a function of donor age. **D**. Probability for a clone detected at some timepoint with frequency 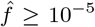 to be detected again at a later time point (with the experimental dataset size). Symbols are empirical estimates. The model predictions show excellent agreement. Error bars in **B.-D**. are propagated from **A**.

Since our approach is probabilistic, it provides as a byproduct the posterior distribution of the fold change of individual clones (see Methods). The average of this posterior over clones agrees very well with the model propagator (4) (Fig. S3), validating its consistency with the data.

The turnover time *τ* increases sharply with age, from a few years at age 21 to ∼ 50 years at age 57 (Fig. 4C). Since the ratio 2*θ/τ* is constrained to be ≈1, this implies that the amplitude of the stochastic stimulations, *θ*^−1^, decreases with age. The TCR repertoire is more dynamic, with faster turnover, for young individuals, who also have a larger rate of introduction of new TCR clones from the thymus than older individuals. At the same time, a turnover time of ∼20 years at the age of 40 suggests that the repertoires of adults remain dynamic despite greatly reduced thymic output.

For individual P6, both TCR*α* and TCR*β* RepSeq samples were available. We recover very similar dynamic parameters for both receptor chains (Fig. 4A-B). This justifies our hypothesis that the bulk sequencing of single chains captures well the dynamics of *αβ* clonotypes.

For comparison, we also applied the naive inference procedure, which allows us to include all 9 patients even when replicates are not available. This inference generally gave much larger rates *τ* ^−1^ and *θ*^−1^ (Fig. S4A), suggesting confounding effects of the noise on both parameters (reversion to the mean for *τ* ^−1^, and larger variance for *θ*^−1^). Indeed, results obtained for a larger valuev of the frequency threshold (*f*_th_ = 10^−4^, Fig. S4B) gave smaller values, and in better agreement with the age dependence.

To ask whether the clonal dynamics depended on the cell type, we separately analyzed the longitudinally sampled CD4 and CD8 repertoires of P6, the only individual for which such data were available. The clone size distribution of CD4 falls off with a larger exponent than that of CD8, meaning that its largest expanded clones are relatively smaller (Fig. S5 A), as already noted [28]. We then applied the naive inference procedure with *f*_th_ = 10^−4^ (since we did not have replicates for the CD4 and CD8 repertoires). The inference (Fig. S5B) reveals that CD4 clones turn over more slowly than CD8 cells (smaller *τ* ^−1^), but also have much smaller fluctuations in their sizes (smaller *θ*^−1^). This result is consistent with a shorter tail of large clones and a larger *α* in CD4 than in CD8 (Fig. S5C).

The inference results can be used to predict the persistence of clones, whose turnover has been discussed in the context of aging [28, 29, 35]. For a given individual, we define persistence as the probability that a clone initially observed at frequency ≤ 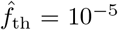 is re-sampled at a later time. This probability strongly depends on the dynamics of turnover of the TCR repertoire, and therefore on the age of the individual, as well as on the time interval between the two samples (Fig. 4D). We can estimate this persistence probability directly from data, and compare it to the predictions of our inferred dynamics, showing excellent agreement. This analysis show that even moderately large clones persist for many years and even decades in older individuals.

Our model assumes that clones have unique trajectories, but that the statistical properties of these trajectories are uniform. However, because of their distinct histories and phenotypical compositions, clones may differ in those dynamical properties. To investigate that possibility, we asked whether the inferred time scales *τ* and *θ* depended on the value of the clone size threshold 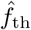. Low values of 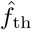 mean that all clones are taken into account in the inference, while high values mean that we focus on the largest clones only. We found that the values of both *τ* ^−1^ and *θ*^−1^ increase with the threshold (Fig. 5A and B) for P3, P4, and P9, suggesting that large clones tend to be more dynamic, with a faster turnover. Therefore, while the overall trends reported in Fig. 4 are still correct, these results imply that the model should be revisited to allow for frequency-dependent dynamics. To measure the frequency dependence of the dynamic parameters more finely, we separately inferred *τ* and *θ* for clones sorted into contiguous intervals according to their initial count 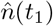. Both time scales showed an approximately linear dependency on the logarithm of the initial frequency (Fig. 5 C and D). This confirms the observation that large clones tend to have faster dynamics than small ones, especially in younger individuals. The two time scales *τ* and *θ* vary in concert, so that their ratio remains approximately constant across frequencies (Fig. S6). As we have argued before, this ratio is linked to the power-law exponent of the distribution of clone sizes at steady state. This exponent can be read off as the slope of that distribution on a double logarithmic scale, which it is consistently observed to be constant in the data (Fig. 1D). Finally, we applied the naive inference procedure to learn the frequency-dependent dynamics of clones in all 9 individuals. As expected, this inference yielded more noisy and less stable results than the full inference, especially at low frequencies for which noise is largest (Fig. S7). The dependence of the inferred parameters on clonal frequency vary across individuals, but confirm a picture in which older individuals have more stable large clones.

**FIG. 5:**
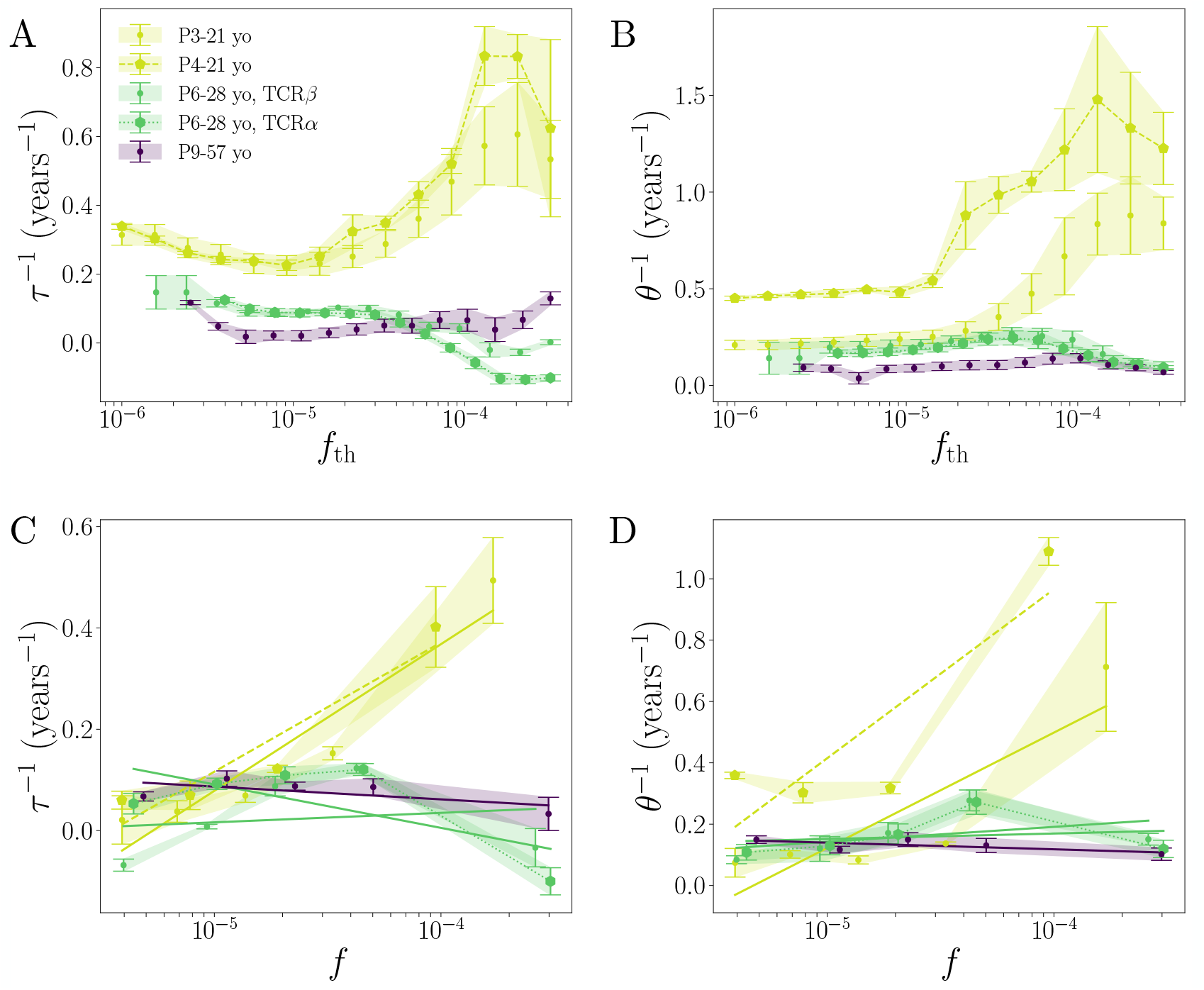
Clonal dynamics are frequency dependent. **A-B**. Results of the full inference as a function of the minimal frequency threshold 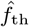 for *τ* ^−1^ and *θ*^−1^. **C-D**. Dynamical parameters as a function of clone frequency. The inference was performed on separate subsets of clones sorted by their frequency in intervals 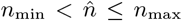, with *n*_min,max_ consecutive numbers in (2, 5, 10, 20, 100, ∞). Error bars are estimated as in Fig. 4.

## III. DISCUSSION

The sizes of T cell clones change constantly throughout the lifetime of an individual, not only due to specific stimulation. We used data sampled on timescales of the order of a year from individuals that did not undergo any strong identified antigenic stimulations to learn the repertoire turnover dynamics. These dynamics include both random unstimulated T cell proliferation and death as well as asymptomatic, or weakly symptomatic antigenic stimulation. We showed that a geometric Brownian motion correctly captures the clone dynamics. This model imposes strict relations that link the exponent of the TCR clone size distribution at steady state, *α*, with the parameters of the dynamics, *τ* and *θ*. We showed that, for all individuals, we were able to predict the measured exponent *α* ≈1 from the inferred dynamical parameters, suggesting that geometric Brownian motion is a good description of the process. Although the actual timescales vary between individuals, with much younger individuals having much faster clone turnover dynamics than older individuals, their ratio is fixed. Indeed, as already noted on a larger cohort of individuals in Ref. [35], the exponent of the power-law distribution of clone sizes does not depend on age.

The source of the faster turnover in younger individuals is not explained by our analysis. It can be linked to a larger thymic output rate [1], imposing a faster turnover. It could also be linked to more a rapid formation of new immune memories at a young age. We did not attempt to separately learn the dynamics of memory and naive pools, since we did not have sorted longitudinal data for which abundance information could be trusted. While it is sometimes assumed that larger clones have a memory phenotype because they must have expanded, a recent study in mice has shown that naive clones can be large as well [14]. It will be interesting to perform a separate analysis of careful sorted naive and memory repertoires in the future using the method described here, especially for individuals of different ages.

More generally, we expect clonal dynamics to be linked to the cellular phenotype, as our preliminary analysis showed for CD4 and CD8 cells. Phenotypes can be characterized with increasing resolution using single-cell expression data [37], which also provides paired TCR information [38]. Future work combining longitudinal sampling with single-cell techniques could help explore the relationship between neutral clonal dynamics and cell type. Additionally, we know that TCR with similar sequences form clusters that often respond to similar stimulants [17, 39], and methods are being developped to annotate repertoire with cluster membership [40] or specificity [41– 45]. As these annotation become comprehensive, one will be able to study the dynamics of specificity clusters, and to assess the persistence of specific immune memories across different immune challenges.

Our current model is based on two effective parameters that describe the timescale for clone turnover, *τ*, and the timescale of random changes, *θ*. Two major assumptions underlie this model. First, it assumes that antigenic stimulation happens repeatedly on short time scales, so that its cumulated effect on longer time scales look like random fluctuations of the net growth rate. Testing this assumption would require longer time traces of the clonal dynamics, to look for memory effects in the clonal growth rates. Second, it assumes that dynamical properties do not depend on the clone size. As observed in Fig. 5, this assumption is only partially verified, with clear violations for 2 of the youngest donors, in which the larger clones display much faster dynamics than the smaller ones. The longidutinal analysis of larger cohorts with a broad age distribution would be required to investigate this effect in detail.

The turnover time scales we infer range from a few years to 50 years, depending on the age of the individual. It has been shown that even sparsely sampled T-cell repertoires can provide a fingerprint that uniquely identifies individuals [46]. The stability of this immune fingerprint is guaranteed for tens of years, provided that the turnover rate is of the order of years or more, as we showed here.

Direct measurements of T cell lifetimes using heavy water [10] give lifetimes of months for memory cells, to a few years for naive cells. These estimates are consistent with our findings: our time scale *τ* is linked to the inverse of the net growth rate of the *clone*, which results from the balance between cell proliferation and death, while experiments based on heavy water measure the turnover of individual *cells*. For instance, memory cells are short lived, but also divide rapidly to compensate for death, so that the size of memory clones remains stable. One may also want to compare our estimate with the previously reported persistence time of clonotypes believed to be of fetal origin, ≈ 37 years [47]. This persistence time is not directly comparable to *τ*, which is the decay rate of the *abundance* of each clone, but it is similar to the characteristic decay of the persistence probability (Fig. 4D), which may be slower. Another caveat is that fetal clonotypes are also primarily naive and take up only a few percent of the repertoire, so that they may not be representative of the overall properties of the clonal dynamics.

Our work was possible because we were able to calibrate the noise using replicate samples. However, replicates are not always available. In this case, the dynamics can still be learned for large clones: we showed using simulations that above a certain frequency threshold, the sampling error becomes small and we can use empirical observations to learn TCR repertoire dynamics directly from read counts. This allows us to correctly estimate the dynamics of large clones without a noise model, if the clones sizes are large at both time points. However, since the repertoire is described by a power law distribution, the role of small clones is far from negligible. An alternative to replicates may be to use close-by timepoints (relative to the time scales of the dynamics) as surrogate replicates. While we had such time points separated by one month for P1, P2 and P7, we did not attempt a full inference on these samples: we did not manage learn a reliable noise model for these donors, because we lacked both the raw sequencing reads and details about the processing procedure (PCR amplification, error correction, etc). In particular, unlike uniquely barcoded cDNA sequencing, PCR amplification of gDNA used for these donors inflates rare clonotypes (as suggested by the low-frequency plateau in the clone size distributions, see Fig. 1D), potentially confounding the analysis.

One of the main conclusions of our work is that repertoires are very dynamic systems, with clone frequencies changing by orders of magnitude on timescales of years, even in the absence of strong known stimulation. This observation challenges our ability to identify responding clonotypes to direct immune stimulation, such as vaccination or diseases. This work builds the ground for inference procedures that not only correct for experimental and biological noise but also for the natural repertoire dynamics. The methods we designed are general and can be used on larger cohorts of individuals presenting different health status, age, and immunodeficiencies features. They provide a promising tool to better understand the maintenance and efficiency of T-cells, enabling to quantify immunosenescence [48], which plays an important role in vaccines performance and cancer research.

## IV. METHODS

### A. Longitudinal data

The datasets analyzed in this study are summarized in Table S1, along with accession number and links to databases.

Data was collected from 4 different studies, which uses two different techniques for repertoire sequencing. Data from [28, 30–32] was generated by sequencing TCR mRNA of PBMCs from healthy individuals, while data from [29] was obtained by directly sequencing genomic DNA (gDNA), as described in detail in each original study.

Briefly, mRNA sequencing was done through cDNA synthesis with template switch allowing for the addition of a unique molecular identifier (UMI), followed by 2-step PCR amplification of the TCR loci (alpha and/or beta), multiplexing, sequencing on an Illumina platform, and processing using the MiXCR software package [49], to obtain lists of clonotypes (V and J segments and Complementarity Determining Region 3 nucleotide sequence) corrected for UMI multiplicity and sequencing errors. gDNA sequencing was done by extracting genomic DNA and performing multiplex PCR to amplify the TCR beta gene before sequencing on an Illumina HiSeq system. Raw data processing was performed using closed software. Since the raw data is not available, we used the processed data provided on the ImmuneAccess platform.

### B. Naive inference

The *naive inference* method directly uses the observed TCR clonal frequencies to learn *τ* and *θ* parameters, assuming that they represent exatly the true frequencies: 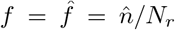. We aim here at maximizing directly the log-likelihood ℒ (*τ, θ*) = log 𝕡 ((*f*_*i*_(*t*_1_), *f*_*i*_(*t*_2_))) | *τ, θ*), which can be expressed by integrating Eq.1:

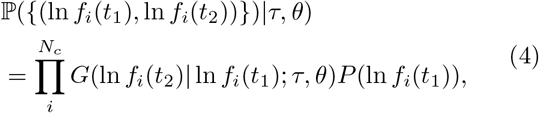

where

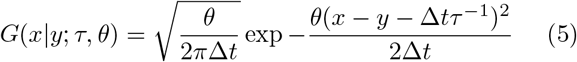

is the propagator of the Brownian motion, Δ*t* = *t*_2_ − *t*_1_ the time interval between the two time points, and where we have assumed that *N*_cell_ is a constant of time. Maximizing the log-likelihood with respect to *τ* and *θ* is equivalent to doing linear regression of ln *f* (*t*_2_) − ln *f* (*t*_1_) against Δ*t*.

### C. Full inference

Using same-day replicates at time *t*_*j*_, we jointly learn the parameters (*α, f*_min_, *a, b*) of the clone-size distribution *ρ*(*f*) = *Cf* ^−1 −*α*^ (for *f*_min_≤ *f* ≤1), and the noise model 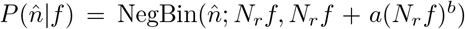 using the NoiSET software [34], where NegBin(*n*; *x, σ*) is a negative binomial of mean *x* and variance *σ*. The learned parameters are reported in Fig. S1.

We then learn parameters of the dynamics by maximizing the likelihood of samples taken at two different time points, using the noise model to account for the discrepancy between true frequencies and sequence counts: For one clone, the full model likelihood reads

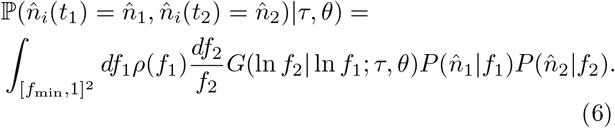

where the noise models are specific to each time point. The maximum likelihood estimator is given by :

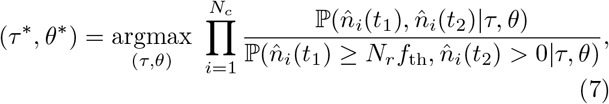

where the denominator accounts for the condition that the clone be included in the analysis: 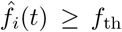 and 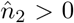. The persistence probability of Fig. 4D is linked to that normalization and is computed as:

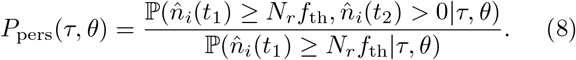

Once the model is learned, the posterior distribution of fold changes *s*_*i*_ ≡ ln *f*_*i*_(*t*_2_) − ln *f*_*i*_(*t*_1_) of each clone *i* is computed through

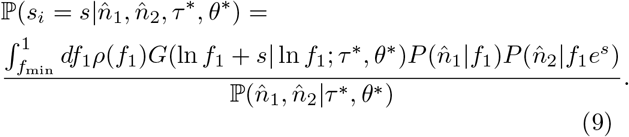

The overall posterior distribution over all clones (solid lines in Fig. S3) is then given by 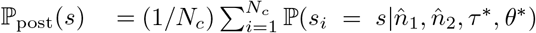. The prior distribution (dashed line), by contrast, is directly given by *G*(ln *f*_1_ + *s*, ln *f*_1_ |*τ* ^∗^, *θ*^∗^), which is independent of *f*_1_. When doing inference in each frequency bin, the product in (7) runs over clones that fall in the bin, and the normalization in the denominator is replaced by the probability to observe 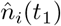 in the bin of interest, and 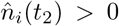. The maximization is performed using the `minimize` function from the Scipy package, with the Sequential Least Squares Programming (SLSQP) method [50] with parameters `tol=1e-8` and `maxiter=300` and initial condition *τ* = 2, *θ* = .5 and constraint *θ*^−1^ *>* 10^−3^.

### D. Synthetic data

Synthetic data was generated by simulating Eq. 1 with a source term producing new clones with rate *S* at initial size *n* = *n*_0_ = 40, and an absorbing boundary condition at *n* = 1. We work with the *x* = ln *n* variable for convenience. The simulation is initialized at steady state, which can be computed analytically [5, 9]. The analytical solution gives us the expected number of cells and clones as a function of the model parameters:

*N*_cell_ = *S*(*n*_0_ −1)*/*(*τ* ^−1^ −*θ*^−1^*/*2), and *N*_*c*_ = *Sτ* ln *n*_0_. Fixing the number of cells to *N*_cell_ = 10^10^, we then compute the number of clones necessary to achieve that size, *N*_*c*_ = *N*_cell_(1 − *τ/*2*θ*) ln *n*_0_*/*(*n*_0_ − 1). We then draw the size 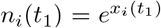 of each clone *i* = 1, …, *N*_*c*_ from the continuous steady state distribution [5]:

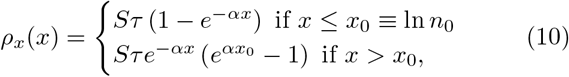

with *α* = 2*θ/τ*.

Then the evolution of each clone from time *t*_1_ to *t*_2_ = *t*_1_ + Δ*t* is determined by the modified propagator with absorbing boundary condition at *x* = 0:

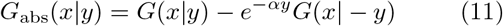

where *G*(*x* |*y*) is defined in (5). In practice, we kill clone *i* with probability 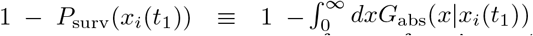, which can be expressed in terms of error functions. Otherwise, its new log-size *x*_*i*_(*t*_2_) is drawn from the distribution *G*_abs_(*x* |*x*_*i*_(*t*_1_))*/P*_surv_(*x*_*i*_(*t*_1_)). In addition, new clones are introduced during Δ*t*. We draw their number from a Poisson distribution of mean *S*Δ*t*, and their introduction times *t* from a uniform distribution in the interval [*t*_1_, *t*_2_]. Then their dynamics is drawn in the same way as for the initial clones, but with Δ*t* = *t*_2_ − *t* instead of *t*_2_ − *t*_1_.

Once the abundances 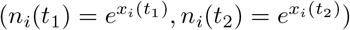 have been determined, the number of reads 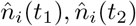 from each time point is drawn from a negative binomial distribution of mean 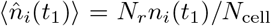 and variance 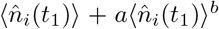, and likewise for 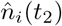, with *N*_*r*_ = 10^6^, *a* = 0.7 and *b* = 1.1.

## E. Code availability

All scripts to produce the figures can be found at https://github.com/statbiophys/Inferring_TCR_repertoire_dynamics/.

## Ackowledgements

This work was partially supported by the European Research Council Consolidator Grant n. 724208 and ANR-19-CE45-0018 “RESP-REP” from the Agence Nationale de la Recherche.

## Supplementary information

**TABLE 1:** Summary of the repertoire samples and individuals used in this study.

| ID | age | sex | # clones | # reads | Tech | DOI ref | data access |
| --- | --- | --- | --- | --- | --- | --- | --- |
| P1 | 18-19 yo | female | $4.08 - 5.11 \cdot 10^5$ | $10.8 - 25.1 \cdot 10^6$ | gDNA | [29] | https://doi.org/10.21417/B7J01X |
| P2 | 18-19 yo | female | $1.55 - 2.93 \cdot 10^5$ | $10.5 - 20.7 \cdot 10^6$ | gDNA | [29] | https://doi.org/10.21417/B7J01X |
| P3 | 21-23 yo | male | $2.04 - 6.44 \cdot 10^5$ | $0.23 - 1.03 \cdot 10^6$ | RNA | [31] | PRJNA493983 |
| P4 | 21-23 yo | male | $1.9 - 10.06 \cdot 10^5$ | $0.28 - 1.79 \cdot 10^6$ | RNA | [31] | PRJNA493983 |
| P5 | 27-30 yo | male | $7.6 - 18.12 \cdot 10^5$ | $1.53 - 6.94 \cdot 10^6$ | RNA | [28] | PRJNA316572 |
| P6 | 28-29 yo | male | $2.62 - 6.38 \cdot 10^5$ | $0.56 - 1.69 \cdot 10^6$ | RNA | [32] | PRJNA577794 |
| P7 | 45-46 yo | female | $1.93 - 2.42 \cdot 10^5$ | $8.86 - 22.8 \cdot 10^6$ | gDNA | [29] | https://doi.org/10.21417/B7J01X |
| P8 | 47-50 yo | male | $1.38 - 9.54 \cdot 10^5$ | $1.53 - 5.93 \cdot 10^6$ | RNA | [28] | PRJNA316572 |
| P9 | 57-58 yo | male | $3.25 - 7.29 \cdot 10^5$ | $0.62 - 1.64 \cdot 10^6$ | RNA | [30] | SRP111073 |

**FIG. S1:**
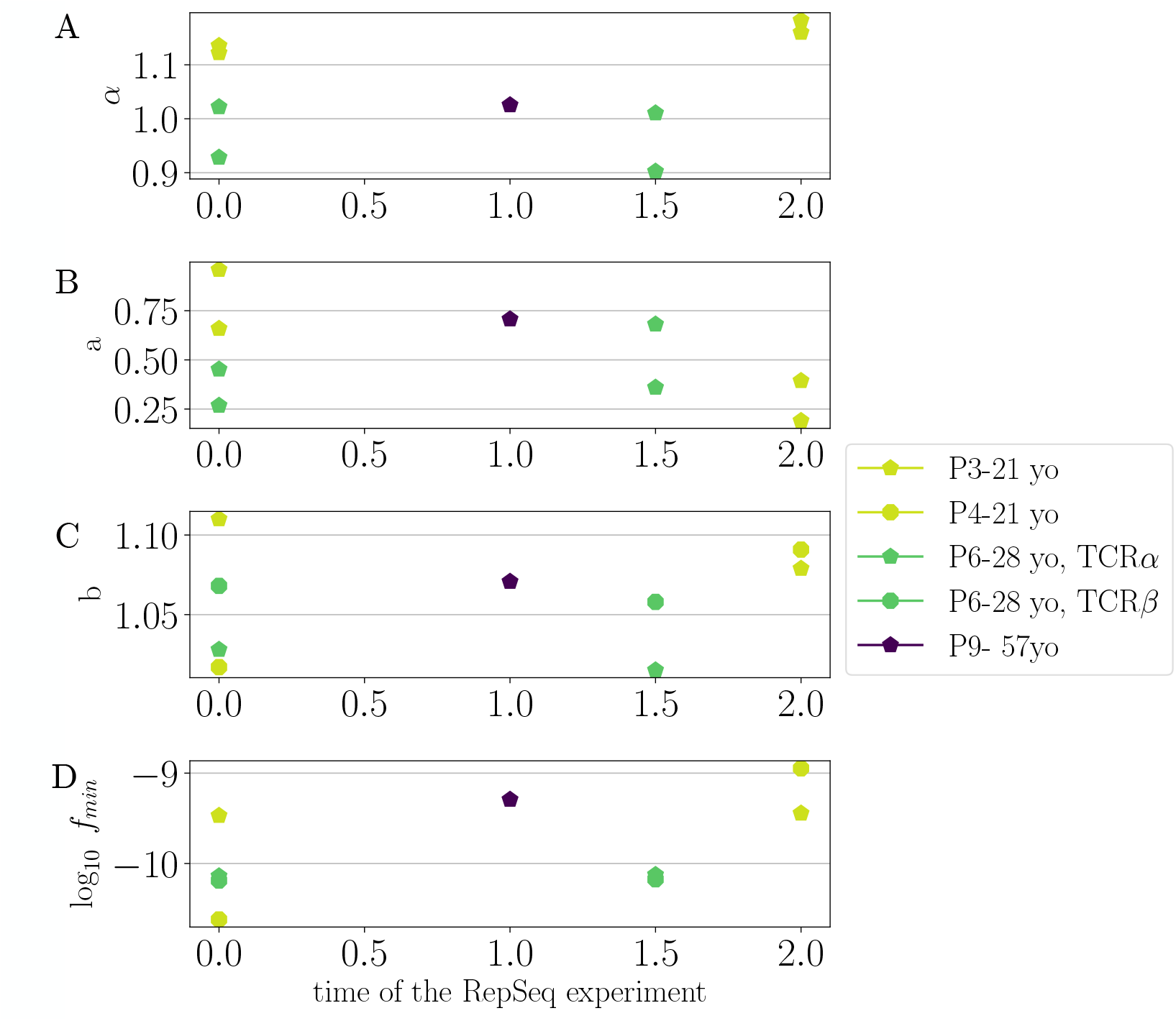
Parameters of the null model, which characterize both the noise model (*a* and *b*) and the prior power law distribution of frequencies (*α,f*_min_). The x-axis represents time in years since the first sample was taken for each individual. Only samples for which duplicates are available are shown.

**FIG. S2:**
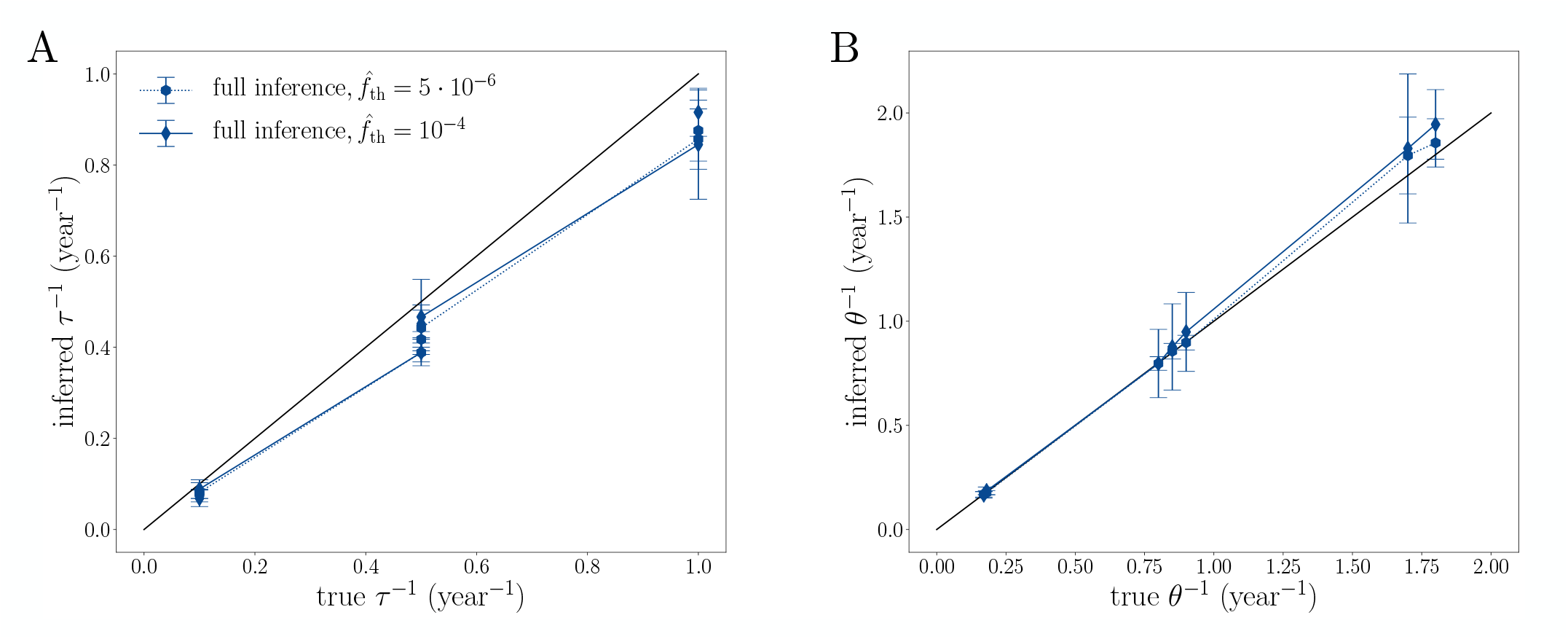
Validation of the inference method for time point with very different number of reads. Parameters: all 9 combinations of *τ* ^−1^ = (0.1, 0.5, 1) year^−1^ and *α* = (1.11, 1.17, 1.25). The number of reads are *N*_*r*_ (*t*_1_) = 10^6^ and *N*_*r*_ (*t*_2_) = 10^5^.

**FIG. S3:**
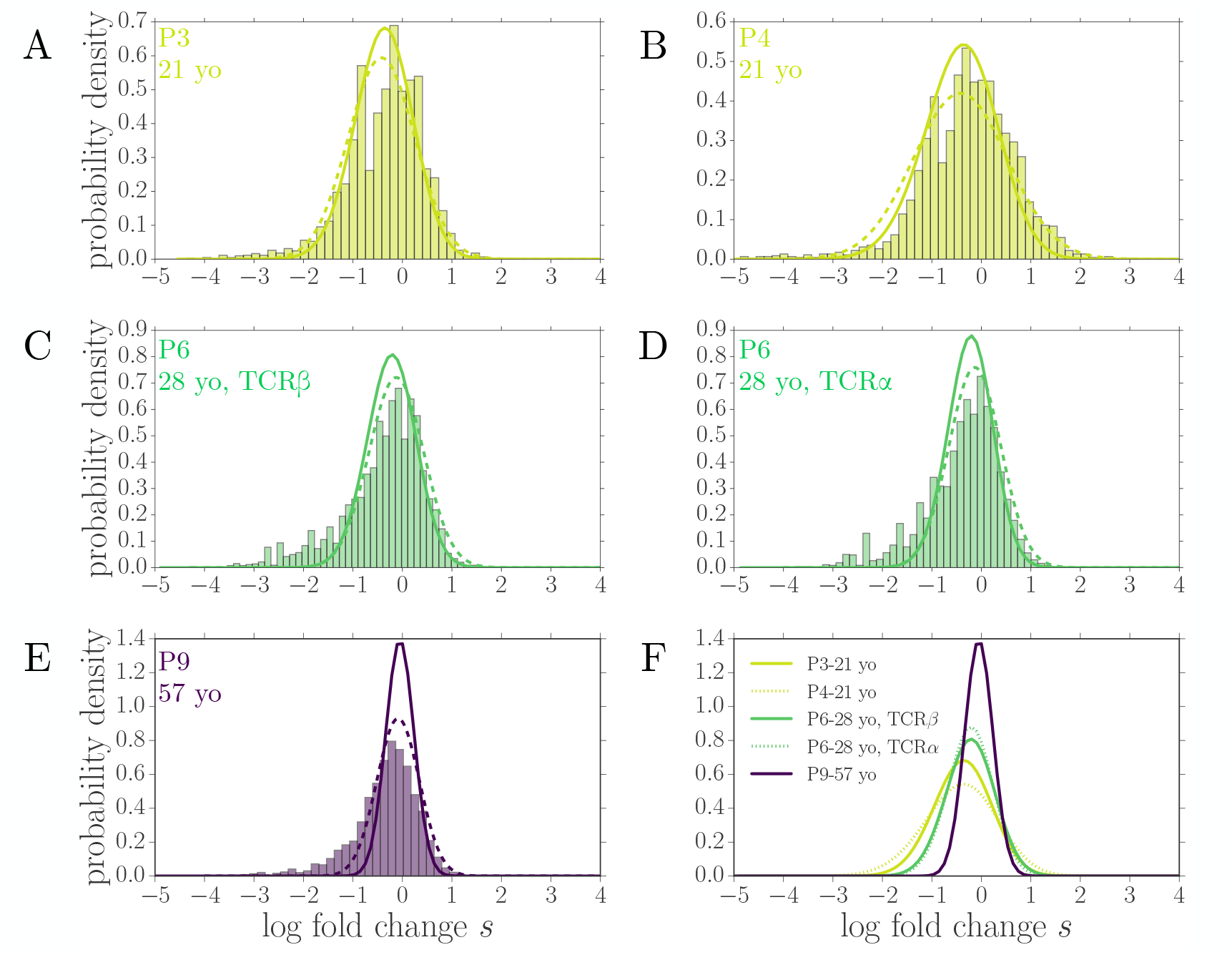
Distribution of log-fold changes *s*. A-E. For each indivudal and chain we compare the naive distribution 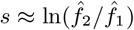 (histogram bars), the prior distribution G(ln *f*1 + *s*, ln f1|τ ∗, θ∗)with the inferred parameters, and the posterior distribution 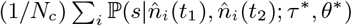. While the naive distribution has an excess of low values due to small number errors, the prior and posterior distribution agree well. **F**. Comparison of posteriors across individuals, showing how both the average decay and its spread decrease with age.

**FIG. S4:**
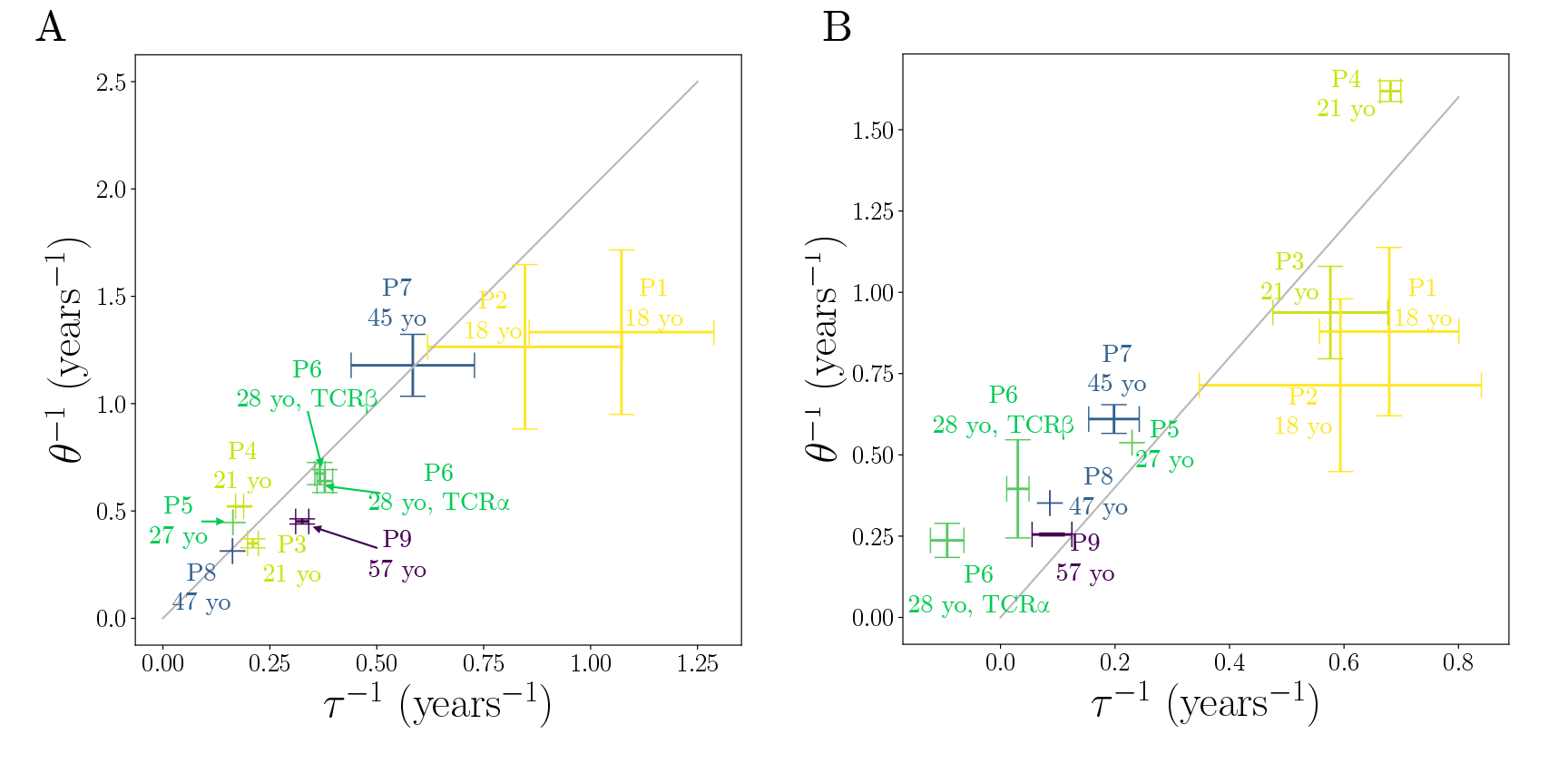
Naive inference of the dynamical parameters on all individuals, with **(A)** *f*_th_ = 10^−5^ and **(B)** *f*_th_ = 10^−4^.

**FIG. S5:**
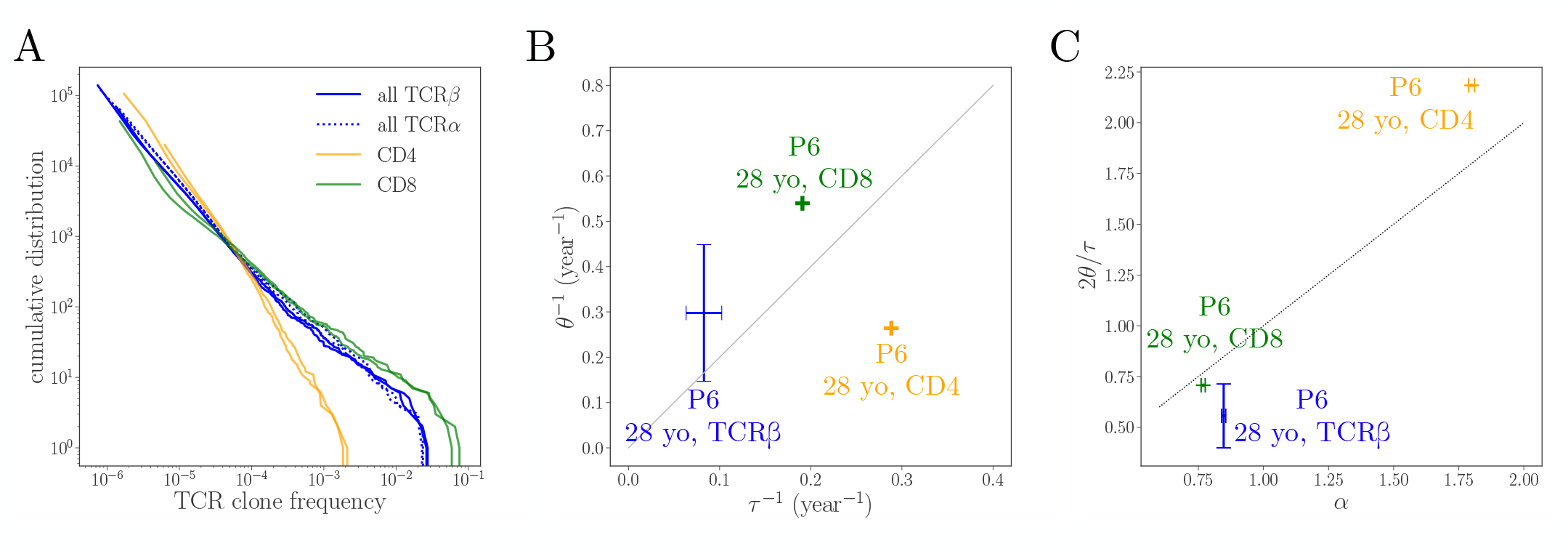
Comparison for the CD4 and CD8 repertoire dynamics in P6. **A**. Clone size distrition in the bulk (alpha and beta chains) and in the CD4 and CD8 beta chain repertoires. The CD4 repertoire has a shorter tail, corresponding to a larger exponent *α*. **B**. Prior and posterior distributions of the log-fold change, as in Fig. S3. **C**. Inferred parameters for each repertoire. **D**. Model prediction (2*θ/τ*) vs measured power-law exponent *α*. The smaller amplitude of frequency fluctuations *θ*^−1^ in the CD4 repertoire is consistent with its shorter tail of large clones.

**FIG. S6:**
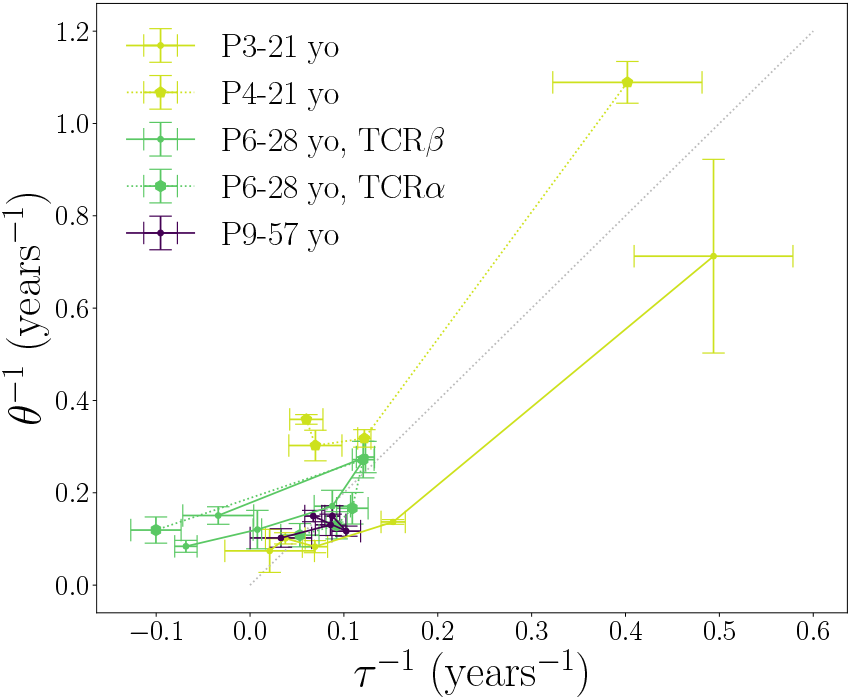
Inferred values of *τ* and *θ* for different individuals and frequencies. Frequency intervals and datapoints are the same as in Fig. 5 C-D.

**FIG. S7:**
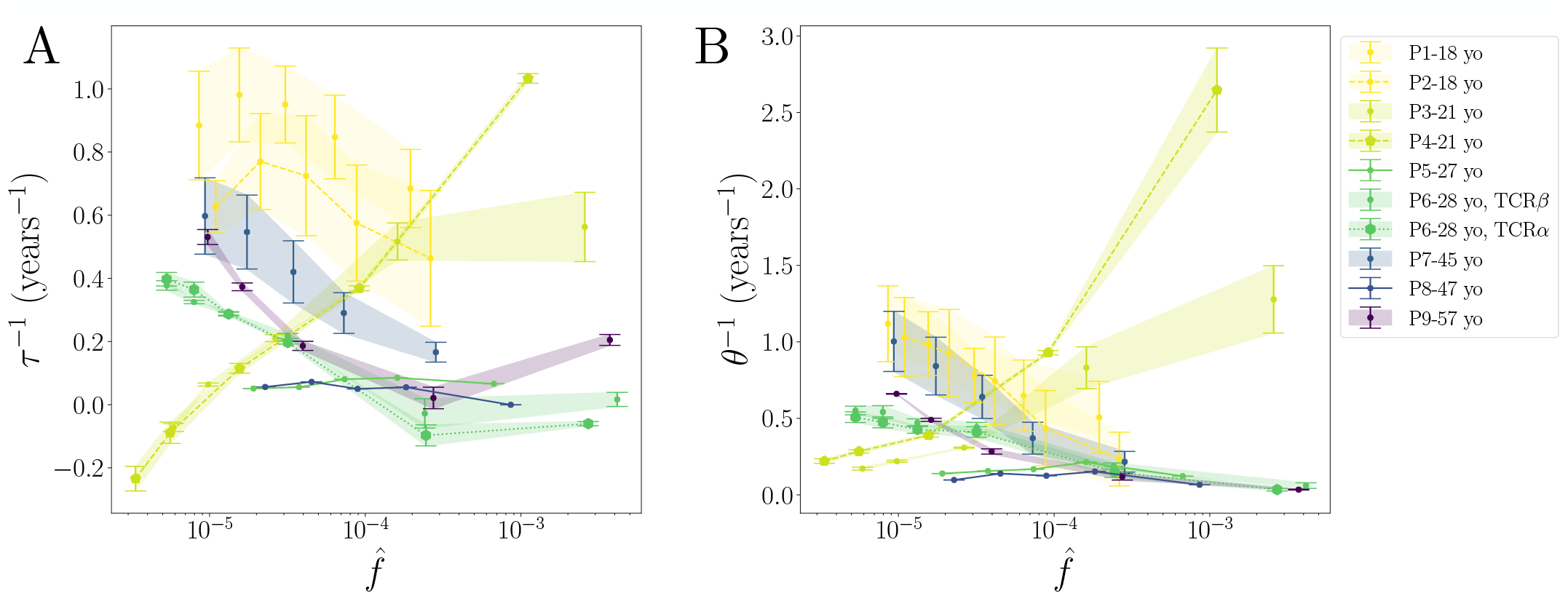
The inference was performed on separate subsets of clones sorted by their frequency in intervals 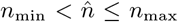, with *n*_min,max_ consecutive numbers in (3, 5, 7, 15, 10, 1000,) for P3, P4, P6, P9, and (100, 200, 400, 800, 2000,) for P1, P2, P5, P7 and P8.

